# Cancer associated fibroblasts mitigate the efficacy of the combination of chemotherapy and BCL-xL targeting in triple negative breast cancer cells

**DOI:** 10.1101/2023.09.26.559287

**Authors:** Lisa Nocquet, Julie Roul, Laurine Duarte, Mario Campone, Philippe P. Juin, Frédérique Souazé

**Affiliations:** Université de Nantes, INSERM, CNRS, CRCINA, F-44000 Nantes, France; Equipe Labellisée LIGUE Contre le Cancer, Paris, France; SIRIC ILIAD, Nantes, Angers, France; ICO René Gauducheau, Saint Herblain, France

**Keywords:** Apoptosis, Breast Cancer, Cancer-Associated Fibroblasts, BCL-2 family, Chemotherapy

## Abstract

Triple negative breast cancers (TNBC) present a poor prognosis primarily due to their resistance to chemotherapy. This resistance is understood to associate with elevated expression of certain antiapoptotic members within the proteins of the BCL-2 family (namely BCL-xL, MCL-1 and BCL-2). These regulate cell death by inhibiting pro-apoptotic protein activation through binding and sequestration and they can be selectively antagonized by BH3 mimetics. Yet the individual influences of BCL-xL, MCL-1, and BCL-2 on the sensitivity of TNBC cells to chemotherapy, and their regulation by cancer-associated fibroblasts (CAFs), major components of the tumor stroma and key contributors to therapy resistance remain to be delineated. By utilizing engineered TNBC cell line MDA-MB-231 deficient in BCL-2 family proteins and BH3 mimetics, we show that BCL-xL and MCL-1 promote cancer cell survival by compensatory mechanisms. This cell line shows limited sensitivity to chemotherapy, in line with the clinical resistance observed in TNBC patients. We elucidate that BCL-xL plays a pivotal role in therapy response, as its depletion or pharmacological inhibition heightened chemotherapy effectiveness. Moreover, BCL-xL expression is associated to chemotherapy resistance in patient-derived tumoroids where its pharmacological inhibition enhances ex vivo response to chemotherapy. In a co-culture model of cancer cells and CAFs, we observe that even in a context where BCL-xL inhibition renders cancer cells more susceptible to chemotherapy, those in contact with CAFs display reduced sensitivity to chemotherapy. Thus CAFs exert a profound pro-survival effect in breast cancer cells, even in a setting highly favoring cell death through combined chemotherapy and inhibition of the main actor of chemoresistance, BCL-xL.

## INTRODUCTION

Triple negative breast cancer (TNBC), characterized by the lack of expression of hormonal receptors and the absence of Erb2 amplification, is associated with a poor prognosis due to its aggressive nature and the absence of effective targeted treatments. Consequently, TNBC patients are often limited to non-targeted chemotherapy, which may not be sufficient to eliminate cancer cells [1,2]. The limited effectiveness of therapy and the development of drug resistance in TNBC is in great part related to pathways regulating mitochondrial outer membrane permeabilization (MOMP). Many anticancer drugs commonly used in clinical practice rely on this mitochondria disrupting event to induce cancer cell death [3]. The BCL-2 family proteins play a crucial role in MOMP, which leads to the release of cytochrome c into the cytoplasm and the activation of a caspase cascade and subsequent cell death through canonical apoptosis or other cell death modes. Cancer cells can survive by altering the balance between anti-apoptotic and pro-apoptotic members of the BCL-2 family [4]. Numerous studies have demonstrated that elevated levels of pro-survival proteins, including BCL-xL, MCL-1, and BCL-2, are frequently observed in cancers and contribute to the tumor escape [5]. BCL-xL and MCL-1 are notably overexpressed in TNBC cells compared to normal counterpart [6–8]. BH3 mimetics, a class of small compounds, antagonize binding of BCL-2 family anti-apoptotic proteins to pro-apoptotic proteins, creating an avenue to offset the excessive expression of these anti-apoptotic proteins [4]. These compounds present an opportunity to gain deeper insights into the exact mechanisms that contribute to chemoresistance. The effectiveness of BH3 mimetics depends on the expression and activity of pro-apoptotic BCL-2 family members and on the balance between the expression of the targeted anti-apoptotic proteins and non-targeted ones [9]. Pro-survival proteins can indeed compensate for each other by taking on released pro-apoptotic proteins induced by BH3 mimetics [4].

The cancer stroma plays a significant role in cancer cell resistance to therapy [10]. CAFs in the tumor microenvironment have been reported to promote resistance to therapy in cancer cells through direct contacts or the local release of soluble factors [11]. We previously showed that in luminal breast cancer, IL-6 secretion by CAFs mitigates the tumor BCL-2 dependency by inducing expression of cancer cell pro-survival protein MCL-1 [12].

In TNBC, identifying the specific pro-survival proteins involved in chemoresistance and investigating whether it is possible to sensitize cancer cells to chemotherapy when CAFs are present by targeting these factors is of particular interest to guide the development of therapeutic strategies. In our study, we used BH3 mimetics and gene silencing techniques to demonstrate the simultaneous and compensatory involvement of MCL-1 and BCL-xL in the survival of the TNBC cell line MDA-MB-231. Furthermore, our findings highlighted the role of the anti-apoptotic protein BCL-xL in mediating resistance to chemotherapy in TNBC. In a co-culture model, we revealed that even when the pro-survival effect of BCL-xL is disrupted, CAFs are still capable of significantly protecting TNBC cells from the effects of chemotherapy. Therefore, our study underscores the need to develop strategies to target CAFs and their interactions with TNBC cells in order to improve the efficacy of chemotherapy in TNBC.

## RESULTS

### BCL-xL and MCL-1 are pivotal anti-apoptotic proteins responsible for the survival of MDA-MB-231 cell line

At first, we assessed the impact of various anti-apoptotic proteins from the BCL-2 family on the survival of MDA-MB-231 cell line. To achieve this, we generated silenced cells for MCL-1, BCL-xL, or BCL-2, designated as sg MCL-1, sg BCL-xL, and sg BCL-2, respectively, using a CRISPR-Cas9 based approach. Our findings reveal that MDA-MB-231 control line express all three anti-apoptotic proteins, albeit with lower levels of BCL-2 (Figure 1A). Western blot analysis corroborated that MDA-MB-231 sg BCL-xL, sg MCL-1, and sg BCL-2 cells exhibit a deficiency in BCL-xL, MCL-1, and BCL-2, respectively (Figure 1A).

**Figure 1.**
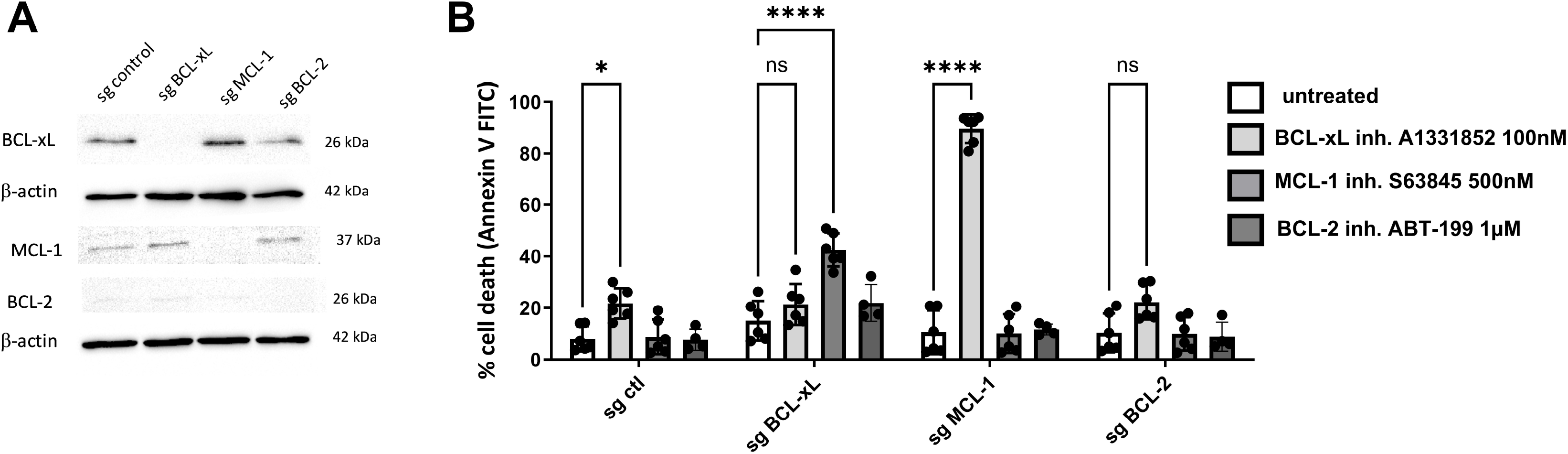
BCL-xL and MCL-1 are pivotal anti-apoptotic proteins responsible for the survival of the MDA-MB-231 cell line. A. Validation of the downregulation of BCL-xL, MCL-1, and BCL-2 expression in engineered MDA-MB-231 cell line through western blot analysis. β-actin expression in the samples serves as a loading control. B. Sensitivity of MDA-MB-231 deficient in MCL-1, BCL-xL or BCL-2 after a 48-hour treatment with BH3 mimetics targeting these proteins (as indicated on the graph) in DMEM containing 0.5% FBS. Cell death is quantified by flow cytometry and the results are expressed as the percentage of cells labelled with annexin-V-FITC. Experiments were conducted in triplicate. P-values were determined by two-factor ANOVA, *p≤ 0.05, ** p≤0.01 and **** p≤0.0001.

Subsequently, we investigated the effects of both loss and pharmacological inhibition of BCL-xL, MCL-1, or BCL-2 on the survival of MDA-MB-231 (Figure 1B). In control cell lines, only the pharmacological inhibition of BCL-xL using A1331852 resulted in cell death in MDA-MB-231. This outcome underscores the critical role of BCL-xL in the fundamental survival of these cells. To explore potential compensatory mechanisms between anti-apoptotic agents when inhibiting only one of them, we employed BH3 mimetics in the different anti-apoptotic-deficient cell lines. Inhibition of BCL-xL with A1331852 in MCL1-deficient cell line led to a substantial 90% cell death, suggesting that simultaneous targeting of both BCL-xL and MCL-1 is necessary to induce massive cell death. Conversely, pharmacological inhibition of MCL-1 by S63845 in BCL-xL-deficient cell line resulted in a less pronounced cell death of only 40%-50%. Inhibition of BCL-2 by ABT-199 in BCL-xL or MCL-1-deficient MDA-MB-231 cells did not induce cell death compared to their respective controls. Similarly, pharmacological inhibition of BCL-xL or MCL-1 in MDA-MB-231 sg BCL-2 cells did not trigger cell death. Hence, BCL-2 does not play a significant role in the survival of MDA-MB-231 cells, consistent with their low expression of BCL-2.

In conclusion, our results highlight that BCL-xL and MCL-1 play significant roles in promoting the survival of MDA-MB-231 cell line. However, the inhibition of either BCL-xL or MCL-1 alone is insufficient to initiate a massive apoptotic response in this studied cancer line. The concomitant inhibition of BCL-xL and MCL-1 clearly increases cancer cells apoptosis, highlighting the presence of compensatory mechanisms between these two proteins.

### BCL-xL significantly contributes to the resistance of the MDA-MB-231 cell line to chemotherapy

One standard of care for triple-negative breast cancers involves the combination of three chemotherapies: anthracycline (such as doxorubicin), alkylating agents (like cisplatin) and anti-metabolites (e.g., 5-fluorouracil). Combination of doxorubicin, cisplatin and 5-fluorouracil resulted in the death of only 25-30% of the MDA-MB-231 cell line as depicted in Figure 2A. To identify the specific anti-apoptotic proteins responsible for impeding chemotherapy effectiveness in these cells, we employed CRISPR-Cas9 gene silencing and BH3 mimetics, as illustrated in Figure 2B and 2C. Combining chemotherapy with the silencing of BCL-xL significantly increased cancer cell death from less than 20% to 60-70%, while silencing MCL-1 or BCL-2 had no substantial impact on sensitivity to chemotherapy (Figure 2B). In accordance with this, combining chemotherapy with pharmacological inhibition of BCL-xL resulted in nearly 100% cell death, whereas MCL-1 inhibition had no effect (Figure 2C). Contrary to BCL-2 silencing, the use of BH3 mimetics ABT-199 in combination with chemotherapy enhanced cell death to 60%, possibly due to an ABT-199 inhibitory effect on BCL-xL at the 1μM dose employed.

**Figure 2.**
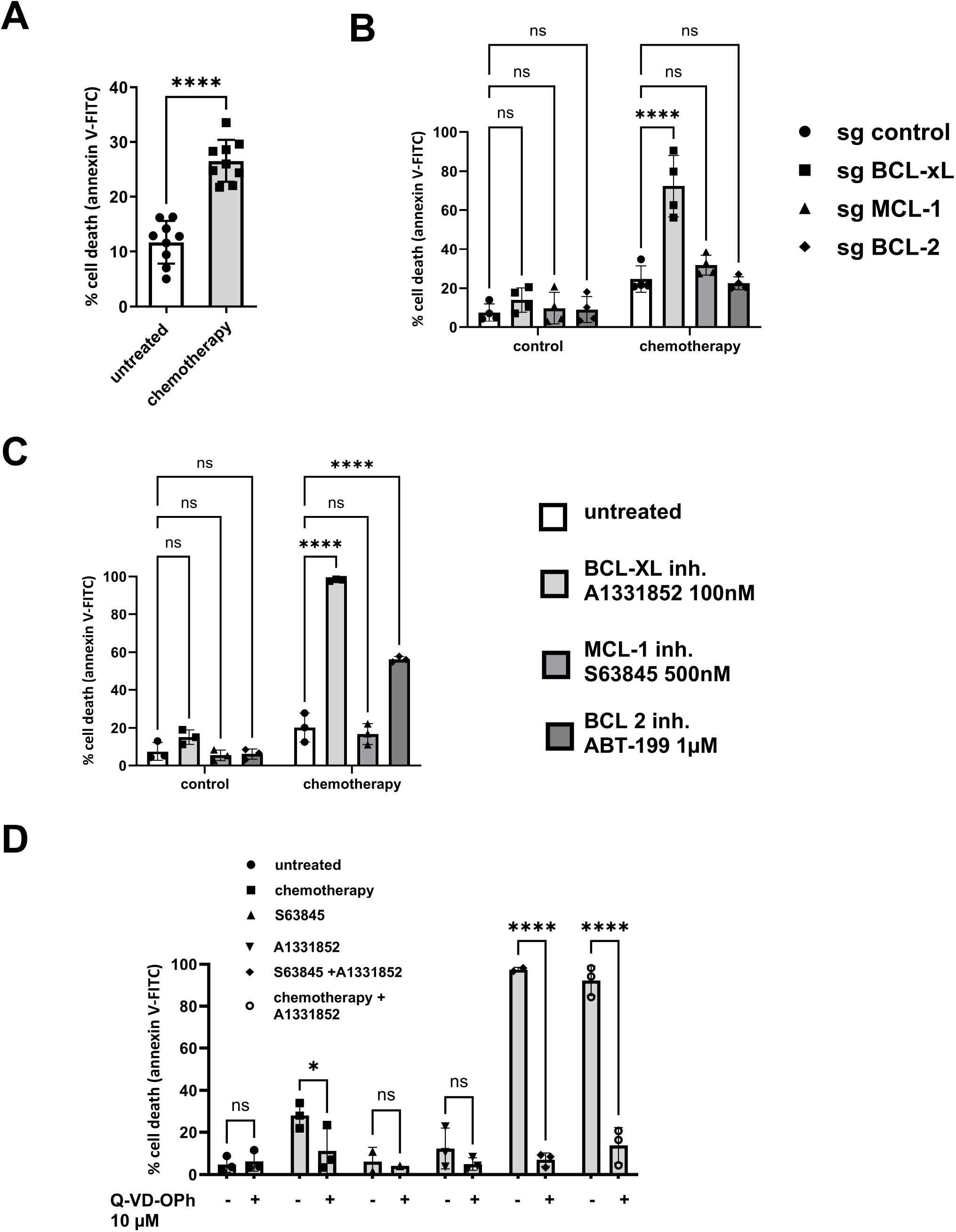
BCL-xL significantly contributes to the resistance of the MDA-MB-231 cell line to chemotherapy. A. Sensitivity of MDA-MB-231 treated 48h with chemotherapy combining Doxorubicin (2.5μM), 5-Fluorouracil (55μm) and Cisplatin (27.5μM). B. Sensitivity of the MDA-MB-231 cells silenced for anti-apoptotic proteins MCL-1, BCL-xL or BCL-2 to chemotherapy. C. Sensitivity of MDA-MB-231 cells treated with chemotherapy combined to BH3 mimetics targeting BCL-xL, MCL-1 or BCL-2 (as indicated on the graph). D. Sensitivity of MDA-MB-231 treated with different combinations of BH3 mimetics and chemotherapy in presence or not of Q-VD-OPh 10μM. Cell death is quantified by flow cytometry. The results are expressed as the percentage of cells labelled with annexin-V-FITC. Experiments were conducted in triplicate. P-values were determined by two-factor ANOVA, *p≤ 0.05 and **** p≤0.0001.

To confirm the apoptotic nature of cell death induced by chemotherapy and BCL-xL inhibitor, we employed a pan-caspase inhibitor (Q-VD-OPh), as shown in Figure 2D. In line with expectations, massive cell death induced by the combination of BH3 mimetics S63845 and A1331852 targeting MCL-1 and BCL-xL respectively was entirely blocked by Q-VD-OPh. Cell death induced by chemotherapy alone or in combination with BCL-xL inhibitor was reversed by Q-VD-OPh. This observation corroborates that cell death induced by chemotherapy, either alone or in combination with BCL-xL inhibitor, is apoptotic in nature.

In conclusion, BCL-xL plays a critical role in limiting the effectiveness of chemotherapy in the MDA-MB-231 cell line.

### CAFs protect MDA-MB-231 cells from apoptosis triggered by simultaneous chemotherapy and BCL-xL silencing

Beyond their intrinsic resistance to chemotherapy, cancer cells survival may also be influenced by the surrounding tumor environment. Consequently, we explored how the presence of CAFs affects the susceptibility of MDA-MB-231 cells to apoptosis using a 2D coculture model. Our findings revealed that the presence of CAFs diminishes the apoptosis in MDA-MB-231 cells triggered by the combination of chemotherapeutic agents (Figure 3A). Simultaneously, the response of CAFs to chemotherapy was variable, with CAF mortality ranging from 20% to 60% under chemotherapy conditions, and notably, the presence of MDA-MB-231 cells did not affect the sensitivity of CAFs to chemotherapy (Figure 3B).

**Figure 3.**
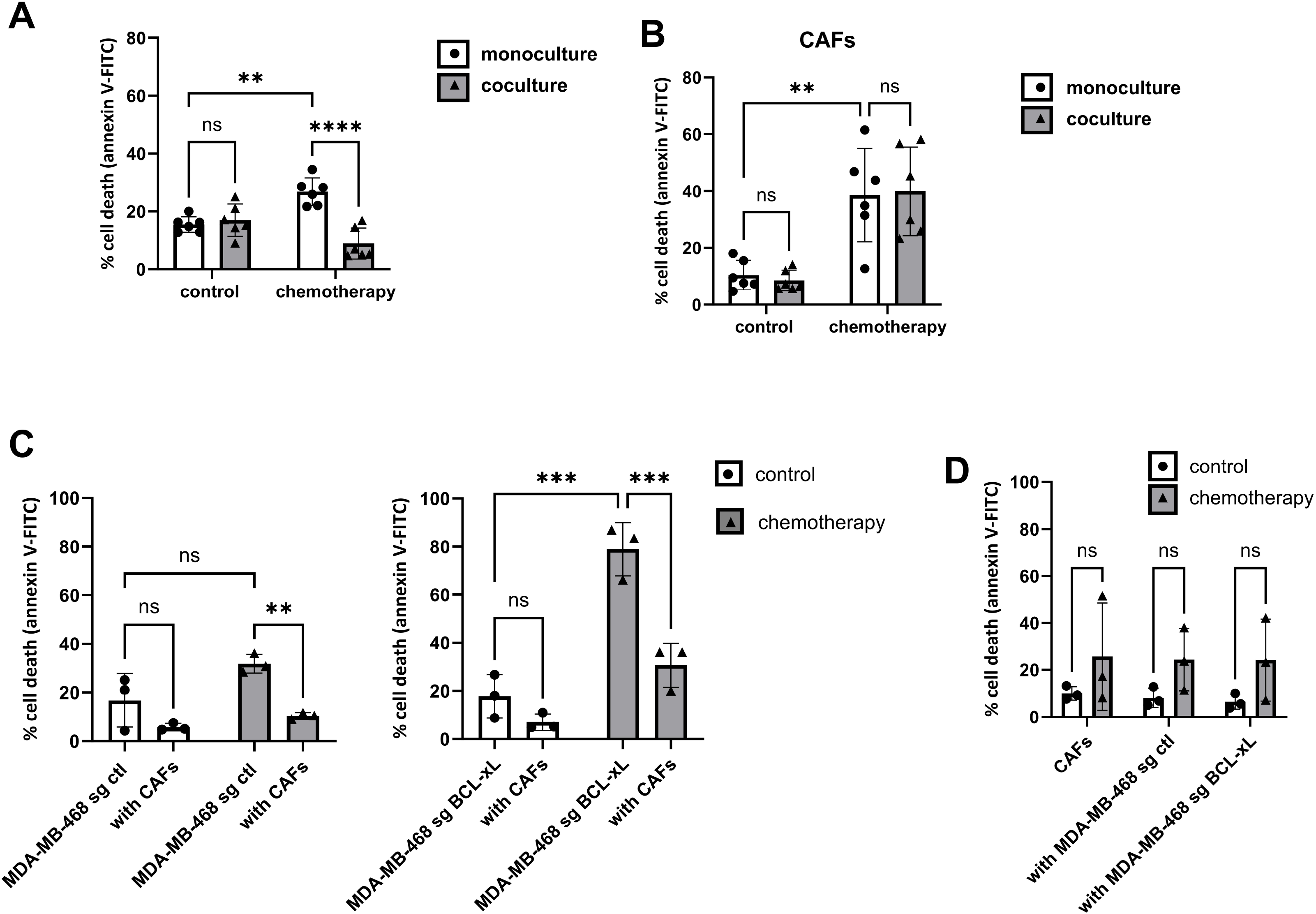
CAFs protect MDA-MB-231 cells from apoptosis triggered by simultaneous chemotherapy and BCL-xL silencing. A. Sensitivity of MDA-MB-231 under chemotherapy in the absence or the presence of CAFs in 2D coculture. B. Sensitivity of corresponding CAFs under chemotherapy in the absence or the presence of MDA-MB-231. C. Sensitivity of MDA-MB-231 control (left) or deficient in BCL-xL (right) to chemotherapy in 2D coculture. D. Sensitivity of corresponding CAFs under chemotherapy in the absence or the presence of MDA-MB-231 control or deficient in BCL-xL Experiments were conducted in triplicate. P-values were determined by two-factor ANOVA, ** p≤0.01, *** p≤0.001 and **** p≤0.0001.

Since inhibiting BCL-xL makes cells more responsive to chemotherapy, we sought to determine whether CAFs could also counteract cell death induced by chemotherapy when BCL-xL was silenced.

As illustrated in Figure 3C, our results demonstrate that the presence of CAFs reverses the cell death triggered by the combination of chemotherapy and BCL-xL silencing. It is worth noting that the survival rate of BCL-xL deficient cells treated with chemotherapy under the influence of stromal pressure closely resembled that of control cancer cells treated with chemotherapy in monoculture (Figure 3C). Concurrently, CAFs displayed variable responses to chemotherapy, and neither the presence of MDA-MB-231 control cells nor MDA-MB-231 cells with BCL-xL silencing influenced the sensitivity of CAFs to chemotherapy (Figure 3D).

Hence, our findings emphasize that CAFs play a protective role against chemotherapy-induced apoptosis, even when the BCL-xL resistance factor is overcome in cancer cells.

### BCL-xL expression is associated with patient-derived tumoroids responsiveness to chemotherapy

To investigate the responsiveness of primary human breast cancer cells to chemotherapy we used 7 breast cancer organoid cultures (hereinafter called tumoroids). Tumoroids derived from treatment naive breast cancer samples were grown *ex vivo* using a stem cell culture method [13] that preserves mixtures of multiple tumor populations of flexible differentiation status [14,15] (see Figure 4A for microscopic analysis). We assessed their response to chemotherapy via FACS analysis and identified variations in sensitivity among the tumoroids (Figure 4B). In particular, tumoroids #2 and #5 showed greater sensitivity to chemotherapy than the other tumoroids, with high sensitivity as early as 0.5μM of chemotherapy. In contrast, tumoroids #4, #6 and #7 were the less sensitive tumoroids to high dose of chemotherapy. Subsequently, we investigated the expression of BCL-xL and MCL-1 in tumoroids using Western blot analysis of bulk lysates (Figure 4C). Remarkably, while MCL-1 expression appears relatively uniform we observed heterogeneity in BCL-xL expression within the tumoroids. Tumoroid #2, #3 and #5 displayed low levels of BCL-xL, while tumoroids #1, #4, # 6 and #7 demonstrated high levels of BCL-xL. Tumoroids expressing the least BCL-xL are the most sensitive to chemotherapy (except tumoroid #3) and conversely the most responsive are those in which BCL-xL expression is high. There is in fact a negative correlation between sensitivity to 0.5 μM chemotherapy and the level of BCL-xL expression in tumoroids (Figure 4D). We treated the most chemoresistant tumoroid (#4) with a combination of chemotherapy and BH3 mimetic antagonist of BCL-xL (A1331852, 100 nM). Consistent with our prior findings in resistant cancer cell line MDA-MB-231, the combination effectively reversed the resistance of tumoroid #4 (Figure 4E). These findings indicate that BCL-xL expression and activity limit the chemosensitivity of patient-derived breast tumoroids. Finally, we observed that the most sensitive tumors (#2 and #5) were rendered more resistant to chemotherapy when treated in the presence of CAFs-conditioned media (Figure 4F). This result confirms the impact of CAFs on cancer cells in tumoroid model of human breast cancer.

**Figure 4.**
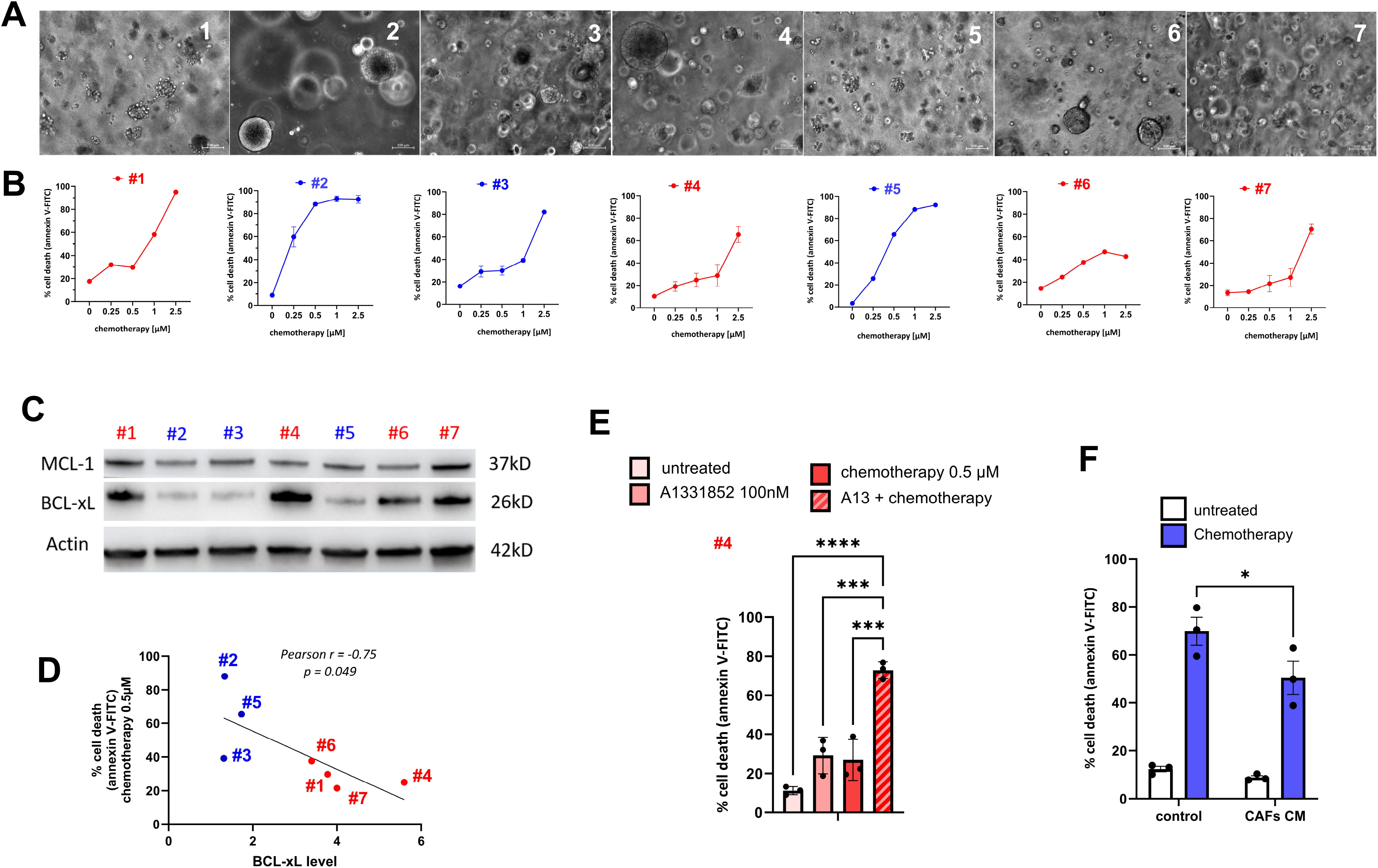
BCL-xL expression affects patient-derived tumoroids responsiveness to chemotherapy. A. Illustration of treatment-naive-patient-derived tumoroids visualized by phase contrast microscopy. B. Heterogeneous sensitivity of patient-derived tumoroids treated with chemotherapy combining Doxorubicin (1X), 5-Fluorouracil (22X) and Cisplatin (11X) for 48h. The results are expressed as the percentage of cells labelled with annexin-V-FITC. Experiments were performed on 7 different tumoroids (#1 to #7). C. Representative experiment of protein expression level of MCL-1 and BCL-xL in patient-derived tumoroids evaluated using western blots. β-actin expression is used as loading control. D. Negative correlation (Pearson correlation coefficient r = −0.75; p value = 0.049) between percentage of cell death upon 0.5μM chemotherapy for 48 hours and BCL-xL levels determined by western-blot (relative to actin) in 7 organoids. E. Sensitivity of tumoroid #4 to treatment of chemotherapy combined with BCL-xL antagonist: A1331852 (100nM). The results are expressed as the percentage of cells labelled with annexin-V-FITC. F. Sensitivity of tumoroid #2 (n=1) and #5 (n=2) to 0.25 and 0.5 μM chemotherapy respectively in the presence or not of CAFs conditioned media. The results are expressed as the percentage of cells labelled with annexin-V-FITC. P-values were determined by two-factor ANOVA, * p≤0.05, *** p≤0.001 and **** p≤0.0001.

## DISCUSSION

In this study, we have demonstrated that triple negative breast cancer cells depend on the presence of BCL-xL and MCL-1 for their survival, which aligns with prior research findings [16–18]. To elaborate further, we investigated the impact of targeting various anti-apoptotic members within the BCL-2 family, namely, BCL-xL, MCL-1, and BCL-2, using a combination of drugs and gene silencing through CRISPR-Cas9 techniques. Our results revealed that the most effective approach for inducing cell death involved simultaneous inhibition of both MCL-1 and BCL-xL. However, it’s worth noting that the outcome varied depending on the method used to target MCL-1 or BCL-xL individually. When we inhibited MCL-1 using the compound S63845 in cells lacking BCL-xL, the level of cell death observed was relatively moderate compared to when we pharmacologically targeted BCL-xL in cells where MCL-1 expression had been genetically removed. It has been previously described that the pro-apoptotic protein BIM shifts from one anti-apoptotic to the other to activate BAX and BAK and engage apoptosis [19]. So, this discrepancy in cell death outcomes could potentially be attributed to differences in the initial binding of the pro-apoptotic protein BIM to either BCL-xL or MCL-1. Our results suggest that a substantial portion of BCL-xL may already be engaged with BIM (or another pro-apoptotic effector), while MCL-1 appears to have a relatively lower binding, possibly being “unoccupied”. This might be explained by the rapid turnover of MCL-1 protein due to its short half-life, and by the fact that its binding by pro-apoptotic effectors fosters its turnover [20]. Binding of BIM to MCL-1 increases under BCL-2 pharmacological inhibition in acute myeloid leukemia cells, thus emphasizing the ability of MCL-1 to take over pro-apoptotics released by other anti-apoptotic proteins [21]. This would underlie the significant role of MCL-1 in promoting “compensatory survival”, in response to an increase of free apoptotic effectors, for instance when BCL-xL is being targeted [17,22].

Currently, aggressive breast cancers, comprising both triple-negative and luminal B subtypes, are primarily managed using conventional chemotherapy regimens that involve a combination of anthracyclines, alkylating agents, and antimetabolites [23,24]. While combination therapies have improved treatment effectiveness in numerous instances, a subset of patients still exhibits resistance. To withstand chemotherapy, cancer cells employ various mechanisms, including inherent resistance to apoptosis governed by members of the BCL-2 family, as well as innate resistance factors like interactions with the extracellular matrix and the secretion of compounds into the tumor microenvironment [25]. We demonstrate here that BCL-xL plays a crucial role in conferring inherent resistance to chemotherapy in cell lines and primary breast cancer cells cultured in tumoroids. This underscores the importance of targeting BCL-xL in breast cancer treatment, especially when used in combination with current therapies. While specific BH3 mimetics designed to target BCL-xL have been developed, they pose a challenge due to on-target platelet toxicity, as platelets rely on BCL-xL for their survival [26,27]. To address this issue, researchers have recently created PROTAC BCL-xL degraders, which recruit E3 ligases that are expressed at lower levels in human platelets compared to various cancer cell lines [28,29]. The approach of co-inhibiting BCL-xL alongside conventional therapies may offer a solution to counter the inherent resistance of cancer cells. However, our findings in MDA-MB-231 cells reveal that even when cancer cells become more responsive to chemotherapy due to BCL-xL inhibition, the presence of stromal CAFs nullifies this effect.

In a prior study, we elucidated that in cases of luminal breast cancer, CAFs possess the ability to diminish the sensitivity of cancer cells to apoptosis. This effect is primarily achieved by promoting the stabilization of MCL-1, mainly through the secretion of IL-6 [12]. The enhanced stabilization of MCL-1 makes these cancer cells more susceptible to MCL-1 targeting, a phenomenon previously observed in other models [21]. In our coculture model, it is possible that additional factors contribute to the resistance of cancer cells to BCL-xL inhibition induced by CAFs. CAFs have been shown to favor chemotherapy resistance in malignant cells via the secretion of growth factors, cytokines and matrix components, which modulate BCL-2 family proteins balance and also effectors of apoptosis downstream mitochondrial permeabilization such as survivin [30]. CAFs could also protect malignant cells from apoptosis by modulating the nutrient context, what has already been shown previously in some studies [30]. Additional investigation is required to determine the specific perturbations that CAFs induce in malignant cells within our model that might account for their protective effects.

In our study, CAFs protect BCL-xL deficient cancer cells against chemotherapy independently of their own sensitivity to treatments, which is heterogeneous. It has already been shown that the growth inhibition effect of conventional chemotherapy on CAFs from breast tumor patients is variable [31]. The maintenance of CAF protective effect despite their disappearance suggests that CAFs could acquire an even more protective effect under chemotherapy. It has indeed already been shown that CAFs induce efficient chemoresistance in colorectal cancer-initiating cells when they are themselves exposed to the same conventional chemotherapy [32]. Our results suggest that the strategy consisting in interrupting the dialog between CAFs and cancer cells could be more suitable than simply eliminating CAFs. It thus appears of particular interest to decipher mechanisms underlying protection from apoptosis induced by CAFs in malignant cells.

Our findings collectively suggest that, even when successfully addressing the inherent chemotherapy resistance mediated by BCL-xL, cancer-associated fibroblasts (CAFs) in the tumor microenvironment will still contribute to the development of acquired chemotherapy resistance. This underscores the need for interventions to overcome this additional obstacle to maximize the effectiveness of therapy.

## MATERIAL AND METHODS

### Primary and cell lines culture

Fresh human mammary samples were obtained from treatment naive patients with breast carcinoma after surgical resection at the Institut de Cancérologie de l’Ouest, Nantes/Angers, France. Informed consent from enrolled patients was obtained as required by the French Committee for The Protection of Human Subjects, and Protocol was approved by Ministère de la Recherche (agreement n°: DC-2012-1598) and by local ethic committee (agreement n°: CB 2012/06).

To isolate CAFs from fresh samples, breast tissues were cut into small pieces in Dulbecco’s Modified Eagle Medium (DMEM Thermo Fisher Scientific) supplemented with 10% FBS, 2mM glutamine and 1% penicillin/streptomycin and placed in a plastic dish [12]. CAFs were isolated by their ability to adhere to plastic. After isolation, the fibroblasts were cultured in the same medium. Fibroblasts were used in the experiments before the ninth passage.

To isolate breast cancer cells from fresh samples, breast tissues were digested by collagenase (20 mg/ml) (Thermo Fisher) for 1h. Breast cancer tumoroids (50 000 cells in 120 μL) were seeded in basement membrane extract (BME, BioTechne) in low adherence 24-well plates (Greiner) and cultured following the procedure previously described by Dekkers et al [33]. Briefly, Advanced DMEM/F12 was supplemented with penicillin/streptomycin, 10⍰mM HEPES, GlutaMAX (adDMEM/F12), 1X B27 (all Thermo Fisher), 1.25 mM *N*-acetyl-l-cysteine (Sigma-Aldrich), 10⍰mM nicotinamide (Sigma-Aldrich), 5⍰μM Y-27632 (AbMole), 5⍰nM Heregulin β-1 (Peprotech), 500⍰nM A83-01 (Tocris), 5⍰ng/ml epidermal growth factor (Myltenyi), 20⍰ng/ml human fibroblast growth factor (FGF)-10 (Peprotech), Noggin (Myltenyi), Rspondin-1 (Miltenyi), 0.1⍰mg/ml primocin (Thermo Fisher), 1⍰μM SB202190 (Sigma-Aldrich) and 5⍰ng/ml FGF-7 (Peprotech). For tumoroids treatment in presence of CAFs conditioned media (CAFs CM), CAFs CM were generated by adding 9 ml of tumoroid medium (Advanced DMEM/F12 as previously described except Y-27632 and A83-01) onto 100mm dishes of 7-12.10^5^ of CAFs. After 48h, CAFs CM were collected, centrifuged (2000g, 5min) and supplemented with Y-27632 and A83-01 before being incubated with cancer cells with treatments for a further 48h before apoptosis analysis.

The human breast cell lines MDA-MB-231 were purchased from American Type Culture Collection (Bethesda, MD, USA) and were cultured in DMEM supplemented with 5% FBS and 2mM glutamine. For the CRISPR Cas9-induced BCL-xL, MCL-1 or BCL-2 gene extinction, single guide (sg) RNA targeting human BCL-xL, MCL-1 or BCL-2 were designed using the CRISPR design tool (http://crispor.tefor.net). The following guide sequences were cloned in the lentiCRISPRV2 vector that was a gift from Feng Zhang (Addgene plasmid #52961): sg BCL-xL: 5’-GCAGACAGCCCCGCGGTGAA-3’, sgMCL-1: 5’-CTGGAGACCTTACGACGGGT-3’ and sg BCL-2: 5’-GAGAACAGGGTACGATAACC-3’. Empty vector was used as control. After transduction, cells were selected using 1μg/ml puromycin and protein extinctions were confirmed by immunoblot analysis. All the cells are tested once a month to ensure they are mycoplasma-free.

### Treatments

Doxorubicin (Selleckhem #S1208), Cisplatin (Selleckhem #S1166) and 5-Fluorouracil (Selleckhem #S1209) were mixed at 2.5μM, 27.5μM and 55μM respectively to obtain chemotherapy treatment. ABT-737 (Selleckhem #S1002), A1331852 (MedChemExpress #HY-19741), ABT-199 (ChimieTek #CT-A199), S63845 (ChemieTek #CT-S63845) and Q-VD-OPh (Sigma-Aldrich #SML0063) were used at indicated concentrations.

### Coculture assays

Cell lines and primary CAFs co-culture was obtained by plating MDA-MB-231 with primary CAFs in monolayer with the ratio 1:3 in DMEM 10% FBS, 2mM glutamine and 1% penicillin/streptomycin. After 24h, the co-culture was maintained in DMEM 1% FBS, 2mM glutamine and 1% penicillin /streptomycin for additional 24h. Treatments were then added as indicated for 48h. Mono-cultures (MDA-MB-231 or CAFs) were performed in the same conditions as controls.

### Apoptosis assay

Cell death was assessed using an Annexin-V FITC binding assay (Miltenyi #130-092-052) performed according to manufacturer’s instructions. Flow-cytometry analysis was performed on Accuri C6 Plus flow cytometer from BD Biosciences. For co-culture apoptosis assay, cell suspension was stained with FITC-conjugated human CD90 antibody (BD Biosciences #555595) prior to Annexin-V APC binding assay (BD Biosciences #550474) according to manufacturer’s instructions.

### Immunoblot analysis

Cells were resuspended in lysis buffer (RIPA, Thermo-Fisher #98901). After sodium dodecyl sulfate– polyacrylamide gel electrophoresis (SDS-PAGE), proteins (20μg per condition) were transferred to a 0.45μM nitrocellulose membrane (Trans-Blot Turbo RTA Transfer kit nitrocellulose midi #1704271) using Trans-Blot Turbo Transfer System (Bio-Rad #1704150, parameters: 25V, 1A, 30 minutes). The membrane was then blocked with 5% non-fat milk TBS 0.05% Tween 20 for 1h at room temperature and incubated with primary antibody overnight at 4°C. The used primary antibodies were: anti-BCL-xL (abcam #ab32370), anti-MCL-1 (Santa Cruz #sc-819), anti-BCL-2 (Dako #M0887) and anti-β-actin (EMD Millipore #MAB1501). The membrane was then incubated with appropriate HRP-conjugated secondary antibody (Jackson Immuno-Research Laboratories, Goat anti-Rabbit #111-035-006 or Goat anti-Mouse #115-035-006) for 1h at room temperature and visualization was made using the Fusion FX from Vilber (Eberhardzell, Deutschland).

### Statistical analysis

Two-way analysis of variance (ANOVA) was used for statistical analysis for overall condition effects with GraphPad Prism 10.2 Software. All data are presented as mean +/-SD of at least three independent experiments.

## Acknowledgements

We thank members of the “Stress adaptation and tumor escape” laboratory for their support. We benefited from technical support from the Cytometry Core facility (CytoCell) of Nantes University. L. Nocquet *was a recipient of French ministry of Higher Education, Research and Innovation* and was supported by a *fellowship from Ligue contre le cancer (CD44)*. This work was supported by INCa (SIRIC ILIAD INCa-DGOS INSERM-ITMO Cancer-18011 and PLBIO021-129NN), and departmental committee of LIGUE Contre le Cancer CD44, CD53 and CD22. P.P.J.’s laboratory is labelized by the Ligue Nationale Contre le Cancer (LNCC). P.P.J. acknowledges the Association Ruban Rose for providing support.

## Author contributions

L.N., J. R., L. D. and F.S. conducted experiments. L.N., P.P.J and F.S. designed the experiments. L.N., P.P.J and F.S. analyzed the data. M.C. gave assistance in collecting tissue samples. L. N., P.P.J. and F.S. wrote the paper. P.P.J. and F.S. obtained funding. P.P.J and F.S. conceived the study and supervised it. All authors contributed to the article and approved the submitted version.

## Ethics Approval and Consent to Participate

Informed consent was obtained from enrolled patients and protocol was approved by Ministère de la Recherche (agreement n°: DC-2012-1598) and by local ethic committee (agreement n°: CB 2012/06).

